# A workflow integrating organ-on-chip culture and correlative 3D light and electron microscopy for microtissue analysis

**DOI:** 10.1101/2024.07.31.605958

**Authors:** Judith M. Schaart, Dorothee Wasserberg, Marcos A. Eufrásio Cruz, Mariska Kea-te Lindert, Robin H.M. van der Meijden, Rona Roverts, Nataliya Debera, Minh Phu Lu, Jeroen Rouwkema, Wouter H. Nijhuis, Andries D. van der Meer, Pascal Jonkheijm, Nico Sommerdijk, Anat Akiva

**Affiliations:** Department of Medical BioSciences, Radboud University Medical Center, Geert Grooteplein Zuid 28, 6525 GA Nijmegen, The Netherlands; Department of Molecules and Materials, Laboratory of Biointerface Chemistry, Organ-on-Chip Centre, TechMed Centre and MESA+ Institute, University of Twente, Drienerlolaan 5, 7522NB Enschede, The Netherlands; Electron Microscopy Center, Radboudumc Technology Center Microscopy, Radboud University Medical Center, Geert Grooteplein Noord 29, 6525 EZ Nijmegen, The Netherlands; Department of Biomechanical Engineering, University of Twente, Drienerlolaan 5, 7522NB Enschede, The Netherlands; Department of Orthopaedic Surgery, University Medical Centre Utrecht, Wilhelmina Children’s Hospital, Lundlaan 6, 3548EA Utrecht, The Netherlands; Department of Bioengineering Technologies, University of Twente, Drienerlolaan 5, 7522NB Enschede, The Netherlands

**Keywords:** organ-on-a-chip, correlave light and electron microscopy, volume electron microscopy, live fluorescence microscopy, bone-on-a-chip

## Abstract

Correlative microscopy approaches offer powerful means to study tissue development across spatial scales, but combining 3D light and electron imaging remains technically challenging. Here, we present a practical workflow that integrates organ-on-a-chip culture with longitudinal fluorescence imaging and volume electron microscopy. By modifying an existing chip platform designed for aligned tissue growth, we demonstrate the feasibility of extended 3D live imaging and subsequent high-pressure freezing of intact microtissues. Fluorescence-guided targeting enables focused ion beam/scanning electron microscopy (FIB/SEM) of selected regions, revealing ultrastructural features such as cellular organization, collagen alignment, and matrix mineralization. While not aimed at new biological discoveries, this study highlights the compatibility and potential of this pipeline for future high-resolution, multiscale studies of tissue morphogenesis and pathology in controlled microenvironments.

## Introduction

In recent years, there has been an increasing focus on replacement, reduction, and refinement in research using animal experiments (the 3R principle).^1^ Therefore, the interest in physiologically relevant cell culture models to study organ development is rising, and much progress has been made.^2,3^ After the establishment of 3D cell cultures, spheroids, and organoids, the organ-on-chip approach was developed to combine microfluidics and 3D tissue engineering to mimic a specific aspect of an organ in a physiologically relevant, dynamic microenvironment.^3-9^ This is usually achieved by the precise control of nutrients and signaling factors in the culture and/or by the application of mechanical stimuli. The highly reproducible organ-on-a-chip devices are regularly used to mimic tissue interfaces, such as the alveolar-capillary interface of the lung^10^ and the intestine.^11^ These types of devices provide great promise for studying the physiological and pathological processes during tissue morphogenesis and for the *in vitro* testing of therapeutic strategies.^3^

A typical design of an organ-on-a-chip device includes a mold-derived transparent material with millimeter-sized chambers in which microtissues can develop. The enclosed, sterile environment of the devices allows for monitoring of the developing tissue for multiple weeks to months using live (fluorescence) microscopy.^2,5^ As the supply of nutrition in the culture environment is highly controlled, the lab-on-a-chip style collection of biochemical cues from the culture medium is also possible.^2^ Additionally, a very attractive feature of microfluidic devices, and in particular organ-on-a-chip systems, is the capability to introduce mechanical stimulation - such as fluid flow or external load - to the growing microtissue to better mimic physiological conditions.

Nevertheless, obtaining high-resolution, ultrastructural information on the developing 3D tissue architecture at specific locations is still challenging. With the current device dimensions, ranging from millimeter to centimeter thickness, high-resolution fluorescence imaging of cellular or intracellular interactions and tissue ultrastructure is limited to the peripheral layers of the culture, as the penetration depth of light microscopes for these types of tissues is limited to about 300 μm. Furthermore, the use of electron microscopy (EM) and particularly volume imaging (vEM),^12,13^ to further investigate the microtissue ultrastructure development in three dimensions is practically unexplored territory for organ-on-a-chip devices. In contrast, for ultrastructure studies of biological tissues, the use of electron microscopy has significantly increased in the last decade, with applications ranging from volume imaging techniques such as array tomography (AT) and focused ion beam/scanning electron microscopy (FIB/SEM) for imaging tissue in the sub-millimeter range or cellular organizations in the micrometer range.^14^ The latest advancement in this field is correlative light and electron microscopy (CLEM),^15^ which uses information from light microscopy to target a specific region of interest for further (advanced) electron microscopy investigation, either of resin-embedded or cryogenic samples.^16^

To allow vEM and CLEM approaches on microtissues in organ-on-chip systems, the typical currently used device dimensions should be reduced, as the current size prohibits structure-preserving EM sample preparation and faithful multimodal sample navigation in correlative imaging approaches.

To close this gap, we have set-up a workflow based on a modified existing chip platform^17^ to perform state-of-the-art imaging for tracking the microtissue development on-a-chip. This microfluidic device serves as a platform for live imaging of microtissues of < 1.5 mm in length, which additionally can be high-pressure frozen for uncompromised (cryogenic) imaging. The demonstrated approach eliminates the need for extensive EM sample preparation that may introduce artifacts into the samples’ ultrastructure and allows the application of correlative fluorescence and electron microscopy.

To demonstrate the utility of our workflow, we used human mesenchymal stem cells (hMSCs) to develop an osteogenic on-chip culture as a first step towards forming a bone-on-a-chip. Compared to other tissues,^18^ the development of advanced, physiologically relevant 3D models for bone, such as bone organoids and bone-on-a-chip devices that include nutrient control, mechanical stimuli, and a hierarchical matrix organization resembling the bone matrix organization, is lagging behind.^19-27^ With the on-chip system presented here, we could use live fluorescence microscopy to monitor the early phases of matrix formation during osteogenic matrix development, a process that extends over multiple weeks.^28^ Subsequent high-pressure freezing of the entire 3D culture and the application of CLEM allowed us to obtain ultrastructural information at precisely defined locations in this complex hierarchical system.

By including fluid flow as a mechanical stimulus, we set a first step towards designing a culture in which early bone matrix formation can be studied in-depth in a physiologically relevant environment. This on-chip system may be further extended by including multiple cell types and by applying compression to resemble the forces sensed during movement to better study the effect of strain on tissue development. Achieving such realistic 3D bone model systems on an advanced imaging platform, where mechanical and (bio)chemical signals can be applied in a dynamic environment, will provide an important tool to enhance the understanding of bone modeling.

## Results

### Design criteria of the organ-on-a-chip system

Bone is a complex tissue^29,30^ in which cell differentiation and the production of a mineralized matrix occur simultaneously. Adding to this complexity are (i) the characteristic aligned organization of collagen observed at multiple length scales,^31,32^ (ii) local differences in the tissue, where on separate locations, different cell types are responsible at the same time for the formation (osteoblasts) and resorption (osteoclasts) of the mineralized matrix, and (iii) the critical influence of mechanical stimuli on bone formation, resorption, and organization.^33-35^

In the past decades, various approaches to generate 3D osteogenic cell cultures have been published,^19,36-38^ but so far none of these has been able to demonstrate all the above features of bone. Only a few 3D osteogenic culture systems have successfully recapitulated the alignment of collagen by allowing matrix-embedded cells to adhere on opposite sides of the cell culture vessel and, through contraction, develop longitudinal stress.^36,39,40^ Details of the collagen organization process have been inferred from high-resolution information recorded at fixed endpoints,^40^ but monitoring this process with high-resolution imaging at defined points in time and space has so far not been possible.

We therefore designed a chip that allows for the first time: 1. to image the whole tissue using fluorescent microscopy with objectives that provide high resolution and the ability to image the whole sample thickness at any given moment, 2. to register sample orientation such that specific regions of interest (ROI) can be monitored over time, 3. to control microtissue dimensions to be small enough for high-pressure freezing as a whole, and 4. targeting an ROI with light microscopy for further in-depth investigation with vEM. Of specific interest for the investigation of bone microtissues are the use of pillars to support the matrix-embedded cells such as to induce the deposition of aligned collagen^36-40^ and the possibility to introduce mechanical stimuli (shear stress, compression) to provide a biomimetic tissue environment.^19^

### An Organ-on-a-Chip Platform for Multiscale Imaging

The above-described organ-on-a-chip device for culturing 3D microtissues was constructed from a polydimethylsiloxane (PDMS) chamber bound to precision cover glasses (Figure 1). To enable fixation of the culture in one place and introduce the possibility of influencing the structural organization of the microtissue, PDMS pillars, stretching throughout the full height of the culture chamber and bonded to the glass coverslip, were introduced in the culture chambers in variable numbers.^17,36,37,39,41-43^ Chamber shapes were linear or rounded, with widths of 1.0 and 2.0 mm respectively (Figure 1A). Chamber heights were kept at 0.4 mm, and longitudinal pilar-to-pillar distances were set at 1.1 mm. These dimensions allowed longitudinal monitoring of the chip with high-resolution fluorescence microscopy (0.4 NA and 1.2 NA objectives), during multiple weeks of cell culture and were compatible with sample preparation for CLEM at specific time points.

**Figure 1:**
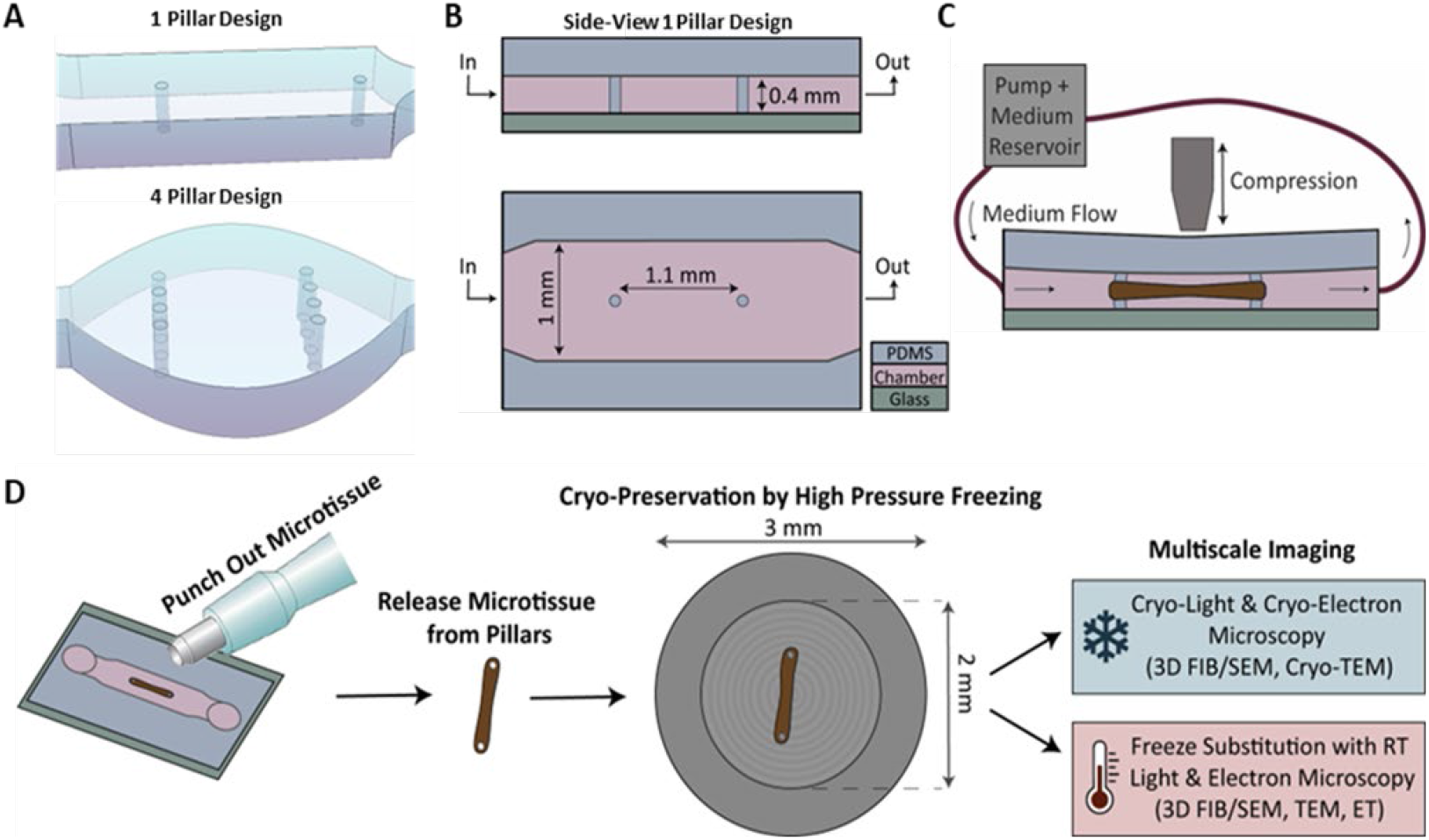
Design of an organ-on-a-chip platform for multiscale imaging. A) 3D representation of two chip designs to influence the structural organization of 3D cell cultures in the chip. PDMS pillars are present in variable orientations to fix the cell culture and modify the culture organization. Designs with microfluidic burst valves were based on Cofiño-Fabres et al.^43^ However, in our design, the PDMS pillars reached the full height of the chamber to fix the position of the pillars. B) Schematic side- and top-view of an organ-on-a-chip platform for fixed and aligned cell cultures. PDMS chips are closed with cover glasses, permitting the use of live fluorescence microscopy. The chamber width is 1 mm in the linear chambers and 2 mm in the round chambers. The height of the culture chamber is 0.4 mm. PDMS pillars of 0.1 mm in diameter are placed at a distance of 1.1 mm to support and fix 3D cell cultures. C) Schematic representation of the application of a continuous medium flow through the in- and outlet of the organ-on-a-chip in which a microtissue is growing between two pillars and the application of compression from the top. D) Schematic overview of the workflow to transfer samples from our organ-on-a-chip platform to light- and electron microscopy: whole samples are punched out of the PDMS chips using a biopsy punch and are released from the pillars. Isolated microtissues are transferred to high-pressure freezing carriers (gray circle) for cryo-preservation by HPF. After cryo-preservation, samples can be used for multiscale imaging by either cryo-light and cryo-electron microscopy or freeze-substitution with subsequent room-temperature light and electron microscopy.

More specifically, to keep the microtissue integrity for EM investigation as close as possible to the live experiment conditions (i.e. minimizing sample preparation artifacts for EM), the chip dimensions were chosen to allow high-pressure freezing (HPF, Figure 1B, D). HPF allows physical fixation of the tissue and replaces conventional chemical fixation to minimize fixation artifacts.^44^ Since, the maximum sample thickness for HPF is 200 μm,^45,46^ the total height of our culture could not exceed this number. The chamber height was set at 400 μm, which allowed the unrestricted inflow of the viscous suspension of cells in their supporting matrix and which, after the expected contraction of the osteogenic cell culture,^40^ resulted in a 3D culture with a height of approximately 100 μm. Furthermore, the carriers we used for HPF have an inner diameter of 2 mm, which defines the limit for the length of the culture (Figure 1B, D). To ensure that the developing microtissues would fit this dimension, the 100 μm wide PDMS pillars that stabilize the culture were placed at a distance of 1.1 mm. This allowed the undisturbed transferring of the whole microtissue to an HPF carrier at any given time point indicated by live high-resolution fluorescence microscopy using a standard skin biopsy punch and subsequently releasing the sample from the pillars (Figure 1D). After HPF, the sample can be used in a workflow applying cryo-light and cryo-electron microscopy in 3D or alternatively the sample can be prepared for imaging at room temperature with freeze-substitution to replace the vitrified ice by a resin.^44^

To mimic physiological environments as close as possible during culture development, the design of our organ-on-a-chip platform included possibilities to apply a continuous fluid flow resembling blood flow using an Ibidi® perfusion system (Figure 1C, Supplementary Figure S1). In addition, we also introduced the possibility of applying mechanical compression through magnetic actuation (Supplementary Figure S1). Together, these functions allow for the optimization of experimental conditions to experiment-specific requirements and offer the opportunity to test different regimes of mechanical stimulation.

### Longitudinal Monitoring of the Bone Microtissue Development

To visualize the early stages of bone matrix development during osteogenic differentiation, human mesenchymal stem cells (hMSCs) were loaded on chips with various pillar configurations as a suspension in fibrin gels, which initially supported the 3D culture. Previous studies showed that uniaxial fixation of fibrin-supported 3D osteogenic cultures, e.g. by pillars or sutures, results in cell alignment upon cellular contraction and the formation of a bone-like aligned collagen matrix.^31,32,36,39,40^ Shortly after seeding, upon stretching and adhesion, the cells contracted the gel around the pillars, yielding microtissues of different shapes (Figure 2A-B). After 3D microtissue assembly, the culture medium was exchanged for an established osteogenic medium to induce cellular differentiation and matrix formation.^47-49^ Visualization of the entire differentiating cultures in 3D (Figure 2A-B, Supplementary Figure S2) showed the dependence of the culture structure on the number and location of the pillars. After seven days of differentiation, maximum intensity projections showed that not only the organization of the cells (red, actin), but also the organization of the deposited collagen (green) had adjusted to the pillar-induced strains that resulted from the cell contraction around the fixed pillars (Figure 2A, Supplementary Figure S2). In each of these bone-on-a-chip devices, the cultures could be differentiated and fluorescently imaged for multiple weeks, which allowed us also to visualize the mineralization of the developing matrix (Figure 2, magenta, Supplementary Figure S2), a crucial step during bone formation. Interestingly, differences in the intensity of the mineral signal were observed between different locations in the cultures, with a generally increased mineralization at the curved regions of the microtissue surrounding the PDMS pillars (Figure 2B). Since the parallel alignment of collagen is a key characteristic of the bone matrix and defines the mechanical properties of bone,^31,32^ we selected the devices in which two single pillars were fixing the cultures (1 pillar design, one pillar on either side of the chip) for further experiments, as these showed the most unidirectional collagen organization.

**Figure 2:**
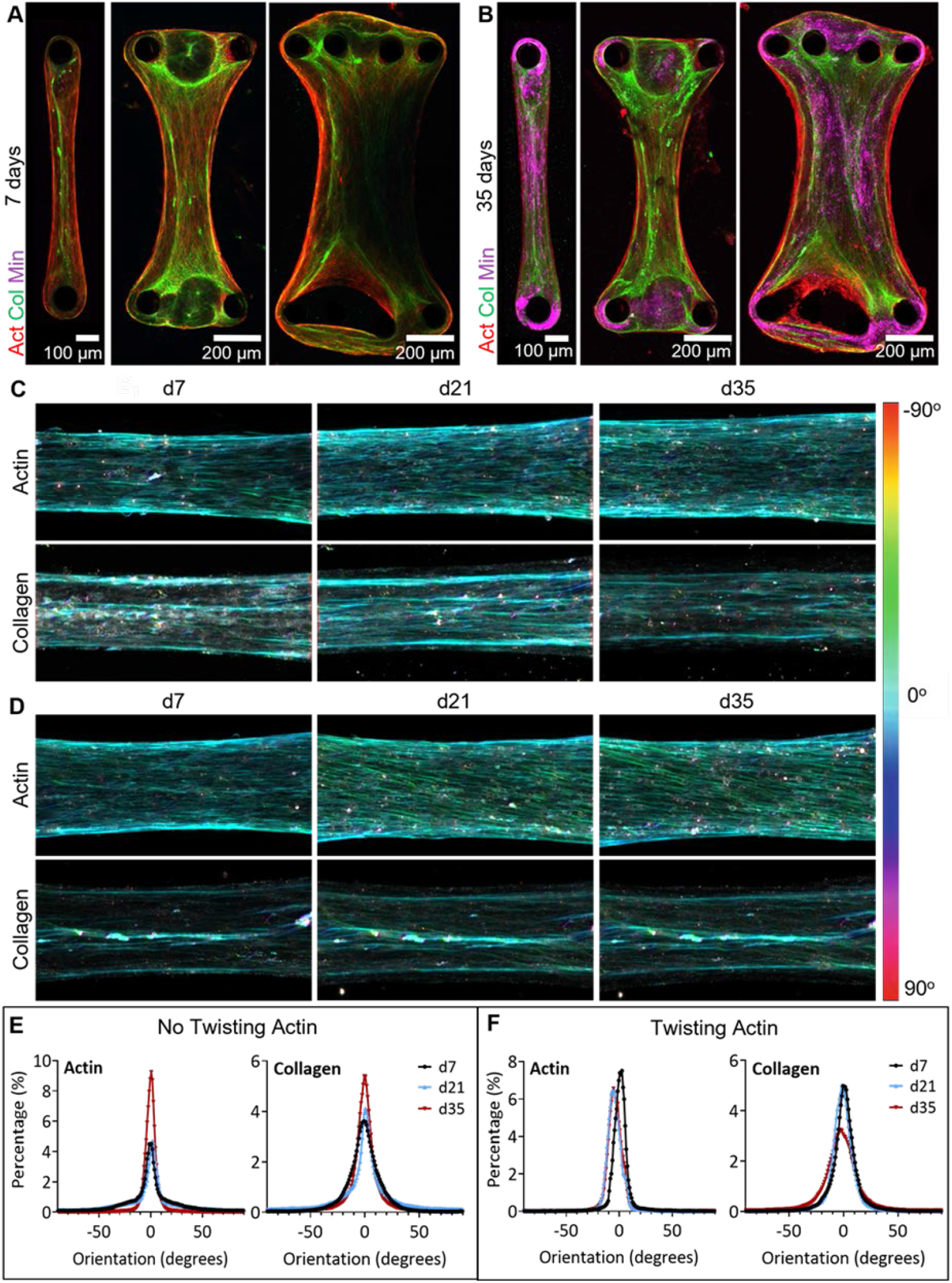
Live monitoring of the development of a bone-on-a-chip culture. A/B) hMSCs were seeded in fibrin gels and differentiated along the osteogenic lineage to grow bone-on-a-chip cultures. Cultures in chips with multiple pillar arrays (1/2/4 pillars per side of the chip) were visualized using fluorescence microscopy after 7 and 35 days of differentiation under static conditions. Maximum intensity projections showed aligned actin (SiR-actin, red) and collagen (CNA35-OG®488, green) patterns after 7 days of differentiation (A). Mineral (Calcein Blue, magenta) was observed mainly in the bent tissue around the PDMS pillars after 35 days of differentiation (B). Corresponding orthogonal views are shown in Supplementary Figure 2A. C-D) Color-based orientation analysis on maximum intensity projections of differentiating cultures (in chips with 1 pillar on each side) showed two types of patterns of actin organization during differentiation: actin orientation remained stable and parallel to the strain direction (C) or started twisting (D) during differentiation. Collagen orientation remained stable over time in both situations. E/F) Orientation histograms showing the preferred orientation confirmed the observation of static (E) or twisting (F) actin during differentiation with a stable collagen orientation. See Supplementary Table S1 for sample numbers.

As the complete microtissues could be imaged live during osteogenic differentiation, weekly monitoring of actin and collagen patterns in the cultures made it possible to study the organizational development of the culture. A color-based orientation analysis of the fluorescence images showed that the cellular actin pattern of the cultures could have two different types of morphologies, which stretched throughout the full culture (Figure 2C-D). The actin signal was either observed in a straight pattern, which approximately aligned to the strain direction between the pillars (Figure 2C), or in a diagonal pattern, in which the actin did not align with the direction of the strain and the cells formed a twisted turn around the culture (Figure 2D, d21/d35).

Interestingly, in samples showing the diagonal pattern, the global actin orientation changed during differentiation, while the collagen orientation, parallel to the strain direction, remained stable over time (Figure 2D). This was also visible in the plotted orientation distributions of the samples (Figure 2E-F), which showed that the preferred orientation of the actin was stable in samples that did not show an actin twist (Figure 2E), while it was changing in samples that showed the twisting actin (Figure 2F). In contrast, the preferred orientation of the collagen remained stable during differentiation in both sample types. This suggests that the local orientation of the collagen fibrils is not dependent on the orientation of the osteoblasts. Based on the imaging of the complete culture over long time scales, we conclude that the observed differences were not dependent on the location of imaging, but the patterns were characteristic of the complete microtissue, which may be related to the pillar geometry in the chamber (Supplementary Figure S4).

### Characterization of the Developing Bone-on-a-Chip in 3D

To better monitor the developmental processes in the cell culture, such as collagen production and mineralization, we analyzed our bone-on-a-chip cultures with advanced live imaging, using Airyscan® fluorescence and Raman microscopy (Figure 3). We first recorded fluorescent z-stack images of the complete culture during osteogenic differentiation to determine a region of interest (Figure 3A, 28 days differentiated). The orthogonal views (Figure 3A_i_-A_ii_) showed that the cultures have a symmetrical shape in the central region, with the most dense actin signal located in the periphery of the culture surrounding a layer of collagen. In the inner core of the culture, cells were also present, even though the density of cells in this region was lower than in the peripheral layer that is in direct contact with the culture medium. Next, we zoomed in to visualize a defined region of interest with high resolution (voxel size: 41 × 41 × 160 nm) using a water immersion objective to minimize refractive index mismatching caused by the various interfaces (glass/medium/culture) that are crossed during imaging (Figure 3B). The maximum intensity projection (Figure 3B, XY) showed the alignment of actin (red) and collagen (green), while the orthogonal views (Figure 3B_i_-B_ii_) showed the organization of the cells and collagen throughout the depth of the culture. The fixation of the culture by the pillars allowed us to identify the orientation of the culture in a reliable fashion, which is essential for this zoom-in approach and the subsequent correlative electron microscopy.

**Figure 3:**
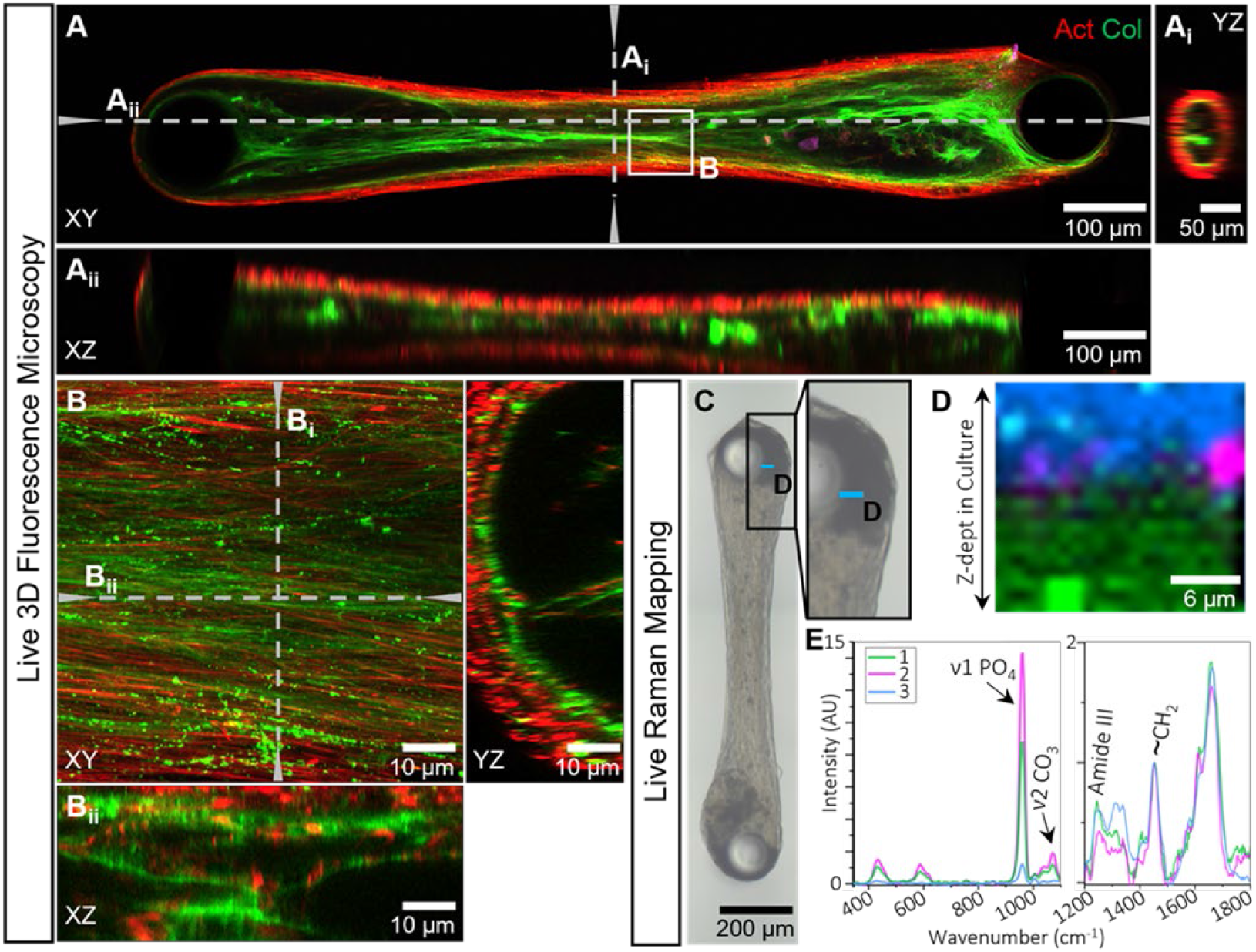
Advanced live imaging of an aligned bone-on-a-chip culture. A-B) Live 3D fluorescence microscopy on-a-chip. A) Live fluorescence microscopy of a complete bone-on-a-chip culture after 28 days of differentiation (A) showed actin (SiR-actin, red) and collagen (CNA-OG®488, green) distributions in 3D. Orthogonal views (Ai: YZ, Aii: XZ) revealed the cylindrical structure of the culture between the two pillars. Orthogonal views were made at the locations indicated by the dotted lines (Ai and Aii). Images were recorded with a 10x objective (NA 0.4, voxel size: 124 × 124 × 1420 nm). B) High-resolution imaging (63x, NA 1.2, voxel size: 41 × 41 × 160 nm) of a region of interest in the center of the culture in 3D showed detailed information regarding the collagen and cellular organization in the culture. Collagen (green) was observed as assembled fibrillar structures (lines) and, presumably, newly secreted matrix (dots). Orthogonal views from the locations indicated by the dotted lines. C-E). Live Raman spectroscopy mapping on-a-chip. C) Optical image of a living bone-on-a-chip culture exposed to fluid flow (0.8 ml/min) taken by a Raman microscope after 21 days of differentiation. The region of interest around the pillar where Raman depth mapping was performed is indicated (blue line, D). D) Spatial distribution of the tissue components, analyzed by True Component Analysis (TCA), identified 1) mineralized matrix (green), 2) carbonated mineral precipitation (magenta) and 3) cellular components with a low amount of mineral (blue). E) the corresponding spectra of the different components shown in D. 1) mineralized matrix, 2) carbonated mineral precipitation, and 3) cellular components with a low amount of mineral.

To characterize the chemical composition of the early mineralizing culture in more detail, Raman spectroscopy mapping was performed on a culture after three weeks of osteogenic differentiation in the presence of fluid flow (0.8 ml/min). These flow-exposed cultures showed similar collagen and mineralization patterns as static cultures, but mineralization was observed earlier during differentiation (day 21, Supplementary Figure S3). Raman depth scans were recorded in two regions, one close to the pillar where the mineral signal was highest in fluorescence microscopy (Figure 3C-E) and one in the center of the culture in which cells and collagen usually showed an aligned morphology (Supplementary Figure S3). True component analysis was used to identify three spectral components in the region close to the pillar, comprising the mineralized matrix (component 1, green), carbonated hydroxyapatite (component 2, magenta), and cellular material with mineral (component 3, blue, Figure 3D-E). The mineralized matrix (component 1) was identified by the presence of the ν1 PO_4_^3-^ vibration at 960 cm^-1^,^50^ together with organic vibrations (amide III (∼1250 cm^-1^), CH_2_ (∼1450 cm^-1^))^50^ and the depth scan showed that this was mainly present near the core of the culture (Figure 3D-E, green). The relative peak heights of the phosphate and organic vibrations show that the matrix in this region of the culture was hyper-mineralized. Component 2 also showed the presence of a clear ν1 PO_4_^3-^ vibration at 960 cm^-1^, as well as a ν2 CO_3_^2-^ at 1072 cm^-1^,^50^ from which we concluded the presence of carbonated hydroxyapatite. From the depth scan it became apparent that this carbonated hydroxyapatite was localized in a round spot, and based on the presence of these two peaks, and the low number of other identified vibrations, we concluded that this was a spot of precipitated mineral (Figure 3D-E, magenta). The top layer of the depth scan (blue, component 3) contained a number of organic peaks (amide III (∼1250 cm^-1^), CH_2_ (∼1450 cm^-1^))^50^ as well as a small mineral peak (960 cm^-1^), which indicated the presence of both cells and mineral in this region. Overall, the distribution of cells and matrix was similar to the distribution observed in fluorescence microscopy. In contrast to the region close to the pillar, the Raman spectra of the region in the center of the culture (Supplementary Figure S3) showed very low mineral-related signals, while the peaks representing cellular material (blue, component 1) and organic matrix (green, component 2) were more pronounced, which was based on the peak around 1340 cm^-1^.^50^ These components were localized similar to the previously observed distribution (Supplementary Figure S3) in which the cells surrounded a collagen layer.

### Correlative Light and Electron Microscopy of a Mineralized Bone-on-a-Chip Culture

After establishing the zoom-in approach to identify and localize a region of interest at multiple magnifications in live imaging, we set up a CLEM experiment to visualize the cell-matrix interactions and assess the mineral deposition during early osteogenic matrix formation in more detail (Figure 4). The applied workflow was similar to the previously reported cryo-correlative workflow of De Beer *et al*.^*51*^ However, instead of identifying the region of interest using the FinderTOP, which can only be used for cryogenic imaging, we now used the sample surface’s unique features caused by the resin embedding (Figure 4A).

**Figure 4:**
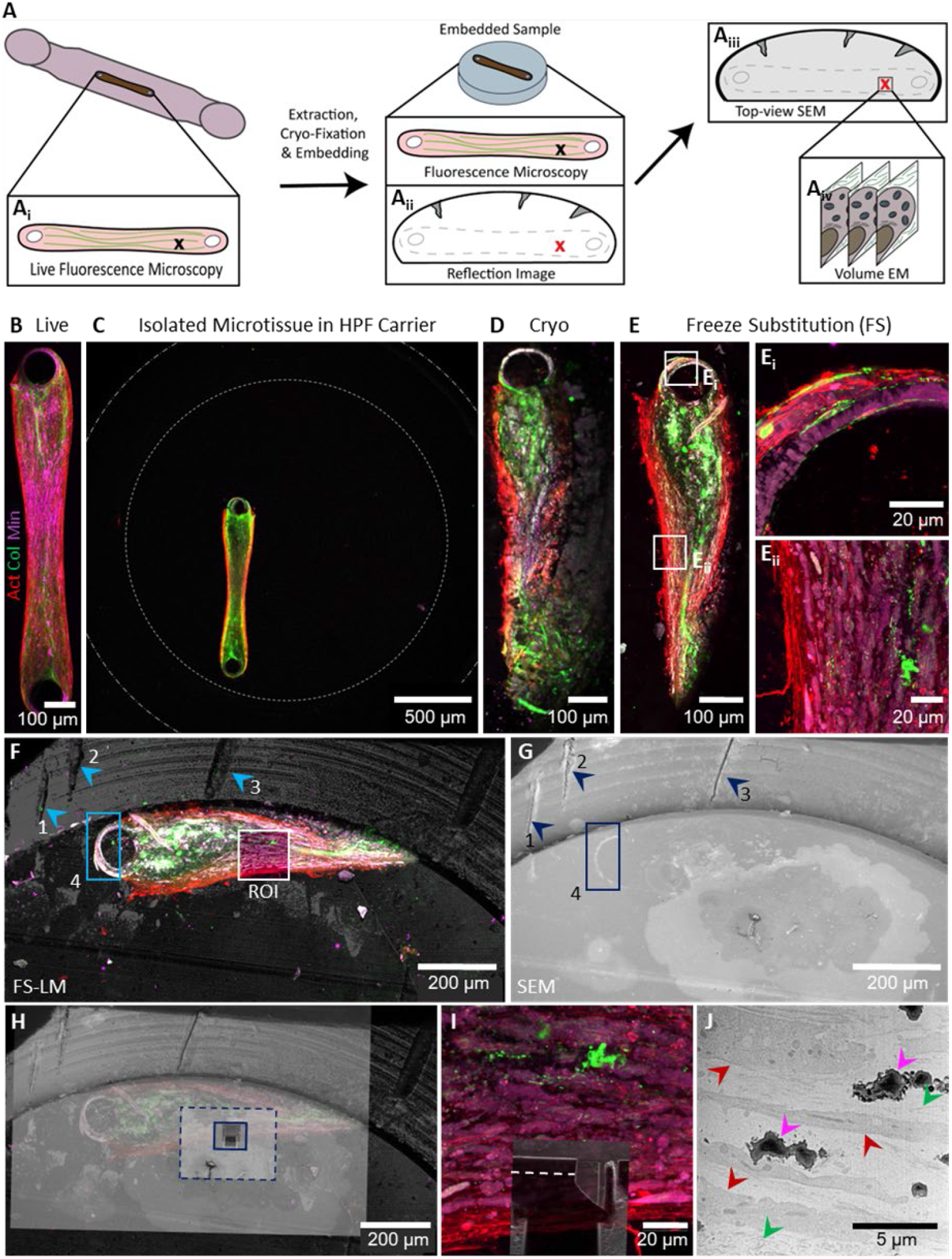
Targeting a bone-on-a-chip culture for multiscale imaging in a defined region of interest. A) Schematic representation of the alignment workflow, starting from live fluorescence microscopy in the culture chamber on chip (A_i_), followed by cryo-fixation and resin embedding. Next, the embedded sample is imaged in parallel with fluorescence and reflection microscopy (A_ii_), after which the sample is transferred to SEM. The SEM top-view (A_iii_) and reflection image are overlayed based on observed irregularities in the resin surface, after which the region of interest can be targeted for volume electron microscopy by 3D FIB/SEM (A_iv_). B) Live fluorescence microscopy of a mineralized bone-on-a-chip culture (actin: SiR-actin, red; collagen: CNA35-OG®488, green; mineral: Calcein Blue, magenta; maximum intensity projection) after 42 days of differentiation, before processing for correlative vEM. C) Overview image of extracted microtissue after transfer to a cryo-fixation carrier for HPF. Dotted lines indicate the inner and outer diameter of the carrier. Image was recorded with a 5x objective (NA 0.2). D) Cryo-fluorescence microscopy confirmed the successful transfer and cryo-preservation of the microtissue (maximum intensity projection). E) Overview of the maximum intensity projection of room temperature fluorescence microscopy after automated freeze substitution (FS) in Lowicryl HM20 resin and determination of regions of interest for high-resolution z-stacks (E_i_/E_ii_). Maximum intensity projection of high-resolution stacks (40x water immersion objective, NA 1.0, voxel size 49 × 49 × 260 nm) around the pillar (Ei) and in the center of the culture (E_ii_). F) Overlay of fluorescence and reflection microscopy of the resin embedded microtissue after freeze substitution for correlation with SEM. Marks for correlation are indicated in blue. G) Top-view of the resin embedded sample in SEM (XY in light microscopy) of the same region as visualized with light microscopy in E, for correlation and identification of the location of the region of interest. Marks for correlation are indicated in blue. H) Overlay of top-view of light and electron microscopy, showing step-wise alignment (dotted blue line) to determine location of the trench for FIB/SEM (solid blue line). I) Overlay of maximum intensity projection of high-resolution fluorescence microscopy of region of interest (E_ii_) overlayed with top-view of trench after FIB/SEM volume imaging has been performed. The location of the FIB/SEM image represented in J is indicated with a dotted line. J) Front-view SEM image from FIB/SEM stack (XZ in light microscopy) in region of interest after 21 μm of milling and imaging. Cells (red arrows), collagen (green arrows), and mineral (magenta arrows) can be identified to analyze the ultrastructure and composition of the sample.

Generally, osteogenic cultures were monitored using fluorescence microscopy for multiple weeks (Figure 4A_i_). At the timepoint of interest, in this case day 42 of differentiation, the microtissue was released from the microfluidic chambers using a punch-out (Figure 1D), imaged again using fluorescence microscopy while it was placed in the HPF carrier, cryo-fixed using high-pressure freezing, and processed for resin embedding by automated freeze substitution to allow subsequent correlative light- and electron microscopy at room temperature. During this process, the shape of the microtissue remained intact, allowing for visual confirmation of the orientation based on the holes in the tissue on the site where the tissue had been surrounding the pillars. After embedding, the

ROI in the resin block was first identified with fluorescence microscopy, after which the reflection imaging mode was used to visualize the surface’s unique features that can also be used for navigating the sample in the EM (Figure 4A_ii_). Next, the fluorescence microscopy data were imported in the electron microscope to allow alignment of the EM view with the optical data (Figure 4A_iii_), after which the location of the ROI was identified and the vEM stack was recorded (Figure 4A_iv_).

The longitudinal live monitoring of actin, collagen, and mineral in the culture (Supplementary Figure S4) again showed that after 35 days of differentiation the first mineral signal was present. The microtissues were kept in culture for one more week to allow the further development of the mineralization until 42 days after the start of differentiation (Figure 4B). One of the microtissues was extracted from the chip within minutes after live fluorescence microscopy to be prepared for electron microscopy. To localize the sample in the carrier, a low-resolution overview fluorescence image of the extracted microtissue was made before HPF (Figure 4C). After HPF, the cryo-preserved sample was visualized again using cryogenic fluorescence microscopy (Figure 4D) to confirm the successful processing of the sample before resin embedding by automated freeze substitution. After embedding, part of the microtissue in the block was imaged using room-temperature fluorescence microscopy to record overviews (Figure 4E) and z-stacks of the ROIs (Figure 4E_i_-E_ii_).

Based on the cellular alignment and mineral signal, region E_ii_ (Figure 4E) was selected as our target region for vEM. To allow for identification of this region in electron microscopy, the reflection image of the freeze-substituted sample was overlayed with the fluorescence imaging of the resin-embedded sample (Figure 4F). The reflection image showed a ring structure with scratch marks imprinted in the resin by the HPF carrier (Figure 4F, blue marks 1, 2 and 3), while the fluorescent image showed the remaining shape of the pillar (Figure 4F, blue mark 4). Next, a top-view image of the sample was recorded with SEM (Figure 4G), where the same ring structures, scratches, and pillar shapes were identified (Figure 4G, blue marks). Using Zeiss Zen 3.7 software with Zen Connect, the three imaging modalities (reflection, fluorescence, and electron microscopy) were overlayed to identify the region for 3D focused ion beam scanning electron microscopy (3D FIB/SEM, Figure 4H). After 3D FIB/SEM, the exact location of the imaged volume within the fluorescence z-stack could be identified by overlaying the top-views in light and electron microscopy (Figure 4I).

As expected, based on the fluorescence microscopy, the FIB/SEM images showed the presence of aligned cells, as well as fibrillar structures, which we identified as collagen (Figure 4J, Supplementary Figure S4). The presence of aligned cells was observed through the complete stack, and cellular organelles could be recognized (marks in Supplementary Figure 4B). While proceeding with the imaging towards the center of the culture (deeper in the stack), a denser layer of collagen appeared in the bottom region of the field of view (Figure 4J, left upper/lower corner, green arrows). Additionally, electron-dense structures were observed between the cells, which we identified as mineral precipitations that were present throughout the complete imaged volume (Figure 4J, magenta arrows, Supplementary Figure S4).

## Discussion

Organs-on-chip have significant importance for studying the functioning of tissues in a realistic *in vitro* environment. On-chip cultures resembling complex 3D tissues have been developed, such as heart-on-a-chip devices in which mechanical stimulation promotes synchronous beating and improved contractile capability in response to electric pacing.^52^ Next to offering a wide range of custom-made organ-on-a-chip platforms for various organs,^4^ microfluidic culture platforms have also been standardized for commercialization in the past years by several companies, mainly focusing on pharmaceutical applications.^2,8,53^ Surprisingly, so far, these existing organ-on-a-chip approaches do not yet allow high-resolution 3D imaging of the entire culture, nor the application of targeted 3D EM approaches, despite the enormous popularity of volume electron microscopy (vEM) and correlative light and electron microscopy (CLEM).

In this study, we have set-up an imaging workflow using a modified organ-on-a-chip platform^17^ that can be used as an advanced 3D imaging platform. Our workflow enables live fluorescence imaging of the entire developing microtissue without compromising on resolution to visualize local variations in the culture during development, which could be overlooked when regional imaging is performed. Furthermore, our device is uniquely suitable for CLEM applications, which allows us to visualize the microtissue ultrastructure in a targeted region of interest with nanometer detail.

Our organ-on-a-chip device is tailored for multi-scale imaging by optimizing our culture chamber to a minimized confined space in which a small 3D microtissue (around 100 × 100 × 1100 μm) could be grown. These dimensions not only allow for live fluorescence microscopy of the entire sample during tissue development, but also allow artefact free cryo-fixation of the microtissue by HPF.^44^ Analyzing cellular organization in organoids by CLEM using automated freeze substitution and targeted analysis of a region of interest by vEM is a valuable tool to study organ development and disease conditions,^19,54,55^ however, a similar analysis of 3D microtissues in microfluidic platforms has not been previously reported. Moreover, the system also allows to follow up with 3D cryoCLEM keeping the sample in its native hydrated state in all stages of the imaging process.^44,51^ Before cryo-fixation, which is an end-point preservation technique, our system allows for the analysis of intermediate stages of the development process by conventional techniques that allow for live read-outs, such as biochemical characterization of the medium outflow, which maximizes the potential output of these devices.

In the past years, the first bone organoids^19-22^ and bone-on-a-chip cultures^23-27^ were reported, which bear great promise for elucidating the still poorly understood bone tissue development pathways. Currently each of these cultures focuses on mimicking specific characteristics of the human bone, such as the woven bone stage^19^ or osteocyte development.^24^ We used our platform to visualize the early phases of osteogenic differentiation, including aligned matrix formation and mineralization.

The pillar fixation of our osteogenic cultures allowed for cellular alignment upon contraction of the supportive fibrin matrix, comparable to what was shown previously for other cell types,^42,43,56^ but in the case of osteogenic cultures, an essential feature to enable the organization of the produced extracellular collagen matrix. Additionally, the fixation-induced alignment of the cells and the matrix they produced allowed us to influence the organization of the culture by adjusting the pillar arrangements (Figure 2, Supplementary Figure S2). This flexibility increases the potential of our organ-on-a-chip device to study not only osteogenic development but also other tissues with specific needs regarding the orientation of cells and matrix.

Ideally, osteogenic culture models should replicate the cell-cell interactions, the hierarchical organization of the cells, and the development of a mineralized matrix. In particular, the importance of mechanical stimulation on bone cell culture development was shown using 3D cultures in bioreactors^19^ or perfused microfluidic devices.^26,27^ The possibility to apply mechanical stimuli, such as fluid flow or compression, was included in our system, to - as close as possible - mimic human physiological conditions in the tissue.

Mechanical stimulation of microtissues in the chip, mimicking *in vivo* flow and compression, further enhances the physiological relevance of this model system. Initial experimental data, together with computational modeling, showed that compression on the chambers of the chips presented in this study is possible at various strengths using a magnetic compression device (Supplementary Figure S1). The future implementation of a rotating magnetic field would enable compression of microtissues not only in a static but also in a dynamic (e.g. cyclic) fashion, to mimic stimuli as sensed for instance by bones during human movement. ^2,57^

Our results demonstrate that it is essential to image the complete, living microtissue without compromising on the spatial imaging resolution to identify local differences in the developing tissue during differentiation and to study sample-specific patterns during the microtissue organization. The local differences in mineralization between curved tissue around the PDMS pillars and the center of the culture (Figure 2) may be explained by local strain differences in the microtissues, which have been identified in previous studies,^42,56^ and would confirm the reported beneficial influence of strain on osteogenic differentiation.^58^

The observed diagonal actin patterns proceeded throughout the complete culture and developed over time, similar to what was previously shown for monolayer mouse osteoblast cultures seeded on curved substrates.^59^ Previous studies also showed that surface curvature was underlying the development of twisted actin patterns by osteoblasts.^59-61^ However, since our cultures all started as comparable, rectangular shapes following the chamber design and subsequently self-organized into tissue straps, no difference in the growth conditions like surface curvature, was expected. Nevertheless, we observed that the actin twisting occurred mainly in chips of which the pillars were slightly bent compared to the pillars in non-twisted samples. We speculate that this results in different strains in the microtissues during development, dependent on the tissue contraction, which could underly the actin pattern formation. Our data show the potential of the presented organ-on-a-chip to study mechanistic processes in tissue development. By systematically adjusting pillar geometry and flexibility in future studies, the presented combination of advanced microscopy with organ-on-a-chip cultures could further elucidate the effect of strain on cell (actin) and collagen organization.

Moreover, our work opens the possibility to extend the analysis of microphysiological devices with targeted ultrastructural analysis in 3D. This for example would allow for the future elucidation of cellular mechanisms involved in extracellular matrix development by specifically focussing on events of interest as observed with the live microscopy, such as the tunnel formation that we previously described in a 3D osteogenic culture.^40^

Beyond osteogenic cultures, our organ-on-a-chip device may be used to study the development of other tissues that show millimeter scale organization, such as muscle,^42,62^ tendon^62^ or cornea,^63,64^ as the chip design could be optimized to stimulate the formation of variable tissue organizations by adjusting the pillar arrangement.^43^ Furthermore, the complexity of the microtissues could be increased by including additional cell types, such as osteoclasts in the bone-on-a-chip culture or endothelial cells for the vascularization of microtissues. Initial studies have already shown the potential of organ-on-a-chip devices for co-culture models.^2,26^ The combination with high-resolution imaging as shown with our platform could further enhance the understanding of the tissue development by providing detailed information on the cell-cell interactions in these co-cultures.

## Conclusions

In the present work, we developed an organ-on-a-chip platform and workflow that enables the detailed and dynamic analysis of the developing microtissues with correlative imaging in cryogenic or room temperature conditions. This device allows for live monitoring of the complete microtissue during multiple weeks of development and the ultrastructural analysis of the sample at a moment and in a region of interest. Furthermore, adaptation of the device to tissue-specific requirements is possible, by modulation of the pillar configurations and inclusion of mechanical cues like fluid flow and compression. As only the cryo-fixation step is destructive for our device, this platform is also suitable to be used in combination with conventional biochemical analysis of the culture medium to monitor secreted factors during tissue development. We therefore expect that the application of this set-up will further improve the current understanding of processes involved in tissue development and structural organization in health and disease, which could help to identify new therapeutic targets for a wide range of diseases, including bone-related diseases, such as osteogenesis imperfecta. Additionally, these platforms are expected to aid in the development of strategies for personalized medicine, by developing patient-specific cultures to test drug efficacy in an individualized, physiologically relevant setting.

## Materials and Methods

Chemicals and cell culture supplements were purchased from Sigma-Aldrich/Merck unless indicated differently. Cell culture media and antibiotics were from Gibco.

### Chip production

A mold for chip production was milled from a 5.5 mm thick, cast Altuglas® poly-methyl methacrylate (PMMA) sheet (Arkema) using a micromill (Datron Neo). After producing the mold, it was cleaned by rinsing with MilliQ, followed by absolute EtOH rinsing and blow drying under a flow of N_2_. Next, dimethylsiloxane monomer of a Dow Sylgard 184 elastomer kit was mixed with the curing agent (ratio 10:1) and carefully degassed *in vacuo* to remove air bubbles. Then, the mold was surrounded with sticky tape to provide a removable barrier and the degassed mixture was poured onto the mold. The covered mold was then degassed *in vacuo* to remove air bubbles and to allow the mixture to completely fill all features of the mold. The degassed mixture on the mold was cured at 80°C for 1 h. Afterwards, the polydimethylsiloxane (PDMS) was carefully peeled off and the unbonded PDMS chips were separated by cutting with a 18 mm wide scraper (Tajima) equipped with a Razar Black Blade®.

Next, 10:1 monomer:curing agent of the Sylgard 184 kit was diluted in heptane (200x). Glass coverslips (#1.5, Thorlabs) were spin coated with the resulting solution (6000 rpm, 30 s, 2000 rpm/s acceleration) and cured at 80°C for 30 min. Using a disposable 1 mm diameter biopsy punch (KAI Medical, B49101), inlets and outlets were punched into the PDMS chips. The unbonded PDMS chips and spin coated coverslips were exposed to oxygen plasma in a Diener Zepto Plasma Cleaner (0.4 mbar, 90 W, 40 s) before bonding and directly after plasma treatment, the PDMS chips were bonded to the coverslips, followed by a short post-curing step (5 min, 70°C) on a heating plate to improve adhesion.

### Cell culture

Human mesenchymal stem cells (hMSCs, passage 2-6) isolated from a healthy female (11 years old) were kindly provided by the department of Orthopaedics of the University Medical Center Utrecht (UMCU). For hMSC isolation, human bone plus marrow waste material was obtained in accordance with the Declaration of Helsinki, with the approval of the Assessment Committee Biobank (TCBio) of the University Medical Center Utrecht (Utrecht, The Netherlands) under the approved number TCBio-18-685, and with the written consent of the parents. Isolated hMSCs were expanded in proliferation medium (PM; αMEM (12571), 10% FCS (F7524), 100 U/mL-100 μg/mL Penicillin/Streptomycin (15140122), 0.28 mM L-ascorbic acid-2-phosphate (AA2P (49752)), 1 ng/mL recombinant human FGF (R&D Systems, 233-GMP-025)) in a humidified incubator (37°C, 5% CO_2_). Upon 80% confluency, cells were split (1:10) by trypsinization (0.05% Trypsin/0.53mM EDTA in PBS) according to standard cell culture protocols.

### Chip seeding and maintenance

Bonded PDMS chips, were coated with Pluronic F-127 (1% in MilliQ, 20 min, RT (P2443)). After coating, the solution was removed and the chips were connected to a vacuum suction system to dry the inside of the channels and culture chambers. Fibrinogen (F8630, 60 mg/mL in αMEM supplemented with 2 mg/mL aminocaproic acid (ACA (A7824)); based on 77% clottable protein) was diluted in complete PM, supplemented with ACA (2 mg/mL) to a final concentration of 6 mg/mL. Thrombin (605157, 50U/mL in αMEM) was diluted in complete PM to a final concentration of 5 U/mL.

hMSCs (passage 3-6) were trypsinized and resuspended in the thrombin solution (2×10^6^ cells/mL). To induce gel formation, the cell/thrombin suspension was mixed with the fibrinogen solution (volume ratio 1:1). Directly after mixing, the chambers of the chips were loaded from one of the inlets. Afterwards, the solution was removed from the channels, and cell suspension only remained in the chambers. The seeded chips (final concentration 1×10^6^ cells/mL) were placed in the incubator (37°C, 5% CO_2_) for 1 h before PM supplemented with ACA (1 mg/mL) was added to the channels, using filter pipette tips as a medium reservoir.

After culturing the chips for two days in PM, the PM was replaced by osteogenic induction medium (OIM, PM supplemented with 6 mM β-Glycerophosphate (βGP, G9422) and 100 nM dexamethasone (D4902)), supplemented with ACA (1 mg/mL). Medium was refreshed 3 times a week during differentiation. To allow degradation of the fibrin scaffold, ACA was left out of the OIM after 14 days of differentiation.

To apply fluid flow, we used an Ibidi® pump system. At the start of the differentiation (day 0), two diagonal inlets of the chip were blocked using sterilized, blunt end needles (19G) filled with cured PDMS. The remaining two inlets were connected via blunt end needles to the Ibidi® Luer connectors of an Ibidi perfusion set (red, 1.6 mm), that were connected to the Ibidi® Fluidic Unit as described by the manufacturer, including pre-run equilibration to remove air bubbles. The syringes connected to the perfusion set each contained 6 mL of OIM (supplemented with ACA (1 mg/mL) during the first 14 days) and the total medium was changed once a week. Flow settings were calibrated to be 0.8 mL/min and with valve switching times of 120 s.

### Live fluorescence imaging of differentiating bone-on-a-chip cultures

Differentiating microtissues were stained live with silicon-rhodamine F-Actin (SiR-Actin, 1 μM + 10 μM Verapamil, Spirochrome, SC001), CNA35-Oregon Green® 488 ^65,66^ (CNA35-OG®488, 0.167 μM TU/Eindhoven, collagen staining) and Calcein Blue (M1255, 0.1 mg/mL, mineral staining). All stainings were diluted in the same medium as the cultures were grown in at that moment. Cells were incubated with the staining solutions for 2 h at 37°C. After staining, the cultures were washed 3x with PBS, and fresh medium was added to the chips. Next, the inlets were closed with a coverglass and the chips were placed upside-down on a object glass. Live cell imaging was performed on a LSM900 upright microscope (Carl Zeiss Microscopy) with Airyscan® 2 detector in multiplex SR-4Y mode, through the cover glass.

For overviews of the cultures a long working distance 5x objective (C Epiplan-Apochromat 5x, NA 0.2, averaging 2x) was used to prepare z-stack images (voxel size: 0.247 × 0.247 × 10 μm). For higher resolution stacks, to allow for orientation analysis, the full culture or regions of interest were visualized using a 10x objective (C Epiplan-Apochromat 10x, NA 0.4, averaging 2x, voxel size: 0.124 × 0.124 × 1.420 μm). To perform high resolution imaging in a region of interest, a 63x water immersion objective (C-Apochromat 63x W.Corr, NA 1.2, averaging 2x, voxel size: 0.041 × 0.041 × 0.160 μm) was used to prepare z-stacks.

### Live Raman microscopy

To prepare the microtissues for live Raman microscopy, the culture medium of unstained, living on-chip cultures was removed and replaced by OIM without phenol red (41061-029). Samples were transferred to the Raman microscope (WITec alpha 300R) and overview optical images were prepared with a 10x objective (NA 0.25). Regions of interest for a depth scan were selected in WITec Control Six (v6.1) software and depth scans were performed using a Zeiss 63x water immersion objective (C-Apochromat 63x W.Corr, NA 1.2) with excitation laser 457 nm at a laser power of 15 mW. Images were recorded with a resolution of 1 × 1 μm with exposure of 4 s per pixel. Recorded spectra were analyzed with WITec Project SIX software. Spectra were baseline corrected with the inbuilt shape correction (radius set at 400) and corrected for cosmic rays. True Component Analysis (TCA) was performed on the sample to identify the different chemical regions, spectra were normalized for the CH_2_ vibration (∼1450 cm^-1^) and color images were created based on the recorded map and selected component.

### Cryo-preservation of microtissues and cryo-fluorescence microscopy

After 42 days of differentiation, the cell cultures were extracted from the chips after live fluorescence microscopy using a skin biopsy punch (diameter 3 mm). The culture was removed from the pillars and transferred to a high pressure freezing carrier (Type A, diameter 3 mm, depth 100 μm, aluminium (M.Wohlwend, 241)) covered in 20% FCS in αMEM (phenol-red free), using a thin needle. The sample was then fluorescently imaged at the LSM900 upright, using a 5x objective (zoom 0.5, single plane), after which the carrier was closed with a flat surface carrier (type B, aluminium (M.Wohlwend, 242)) coated with phosphatidyl choline from egg yolk (1% w/v in ethanol (61755)) and preserved by high pressure freezing (HPM Live μ,^44^ CryoCapCell).

Samples were stored in liquid nitrogen until cryo-fluorescence microscopy. Cryo-fluorescence microscopy was performed on a LSM900 upright with Airyscan® 2 detector, using a Linkam LSM cryo-stage (CMS-196, Linkam Scientific Instruments). The sample was loaded into the cryo-holder (Art. 349559-8100-020, Zeiss cryo accessory kit) which was transferred to a Linkam adapter and imaged under cryogenic conditions to verify the sample location (10x objective, NA 0.4, zoom 0.5, z-steps = 1.42 μm). Afterward, the sample was stored again in liquid nitrogen until freeze substitution.

### Automated Freeze Substitution and fluorescence microscopy of embedded samples

High-pressure frozen samples were placed in pre-cooled FS medium (0.1% Uranyl Acetate in 2.5% (v/v) H_2_O in acetone, -90°C, 72 h) for automated freeze substitution (Leica Microsystems, AFS2). The temperature was gradually increased (3°C/h) to -45°C. After 5 h of incubation at -45°C, the FS medium was replaced by acetone in three sequential acetone incubation steps (10 min), followed by two ethanol incubations (10 min). After this dehydration procedure, a series of Lowicryl HM20 (10%-100%, Table 1) was used for embedding, while gradually increasing the temperature to -25°C (Table 1). The pure Lowicryl HM20 solution was then replaced two more times before fresh Lowicryl HM20 was added and UV irradiation (48 h) was started. Finally, the temperature was increased to 20°C during UV irradiation.

**Table 1:** Lowicryl HM20 Incubation Scheme.

| T<sub>start</sub> | T<sub>end</sub> | Slope | Time | Solution |
| --- | --- | --- | --- | --- |
| -45°C | -45°C | 0°C/h | 6 h | Lowicryl HM20 10% |
| -45°C | -45°C | 0°C/h | 6 h | Lowicryl HM20 25% |
| -45°C | -35°C | 1.67°C/h | 12 h | Lowicryl HM20 50% |
| -35°C | -25°C | 1.67°C/h | 16 h | Lowicryl HM20 75% |
| -25°C | -25°C | 0°C/h | 10 h | Lowicryl HM20 100% |
| -25°C | -25°C | 0°C/h | 10 h | Lowicryl HM20 100% |
| -25°C | -25°C | 0°C/h | 10 h | Lowicryl HM20 100% |
| -25°C | -25°C | 0°C/h | 48 h (+UV) | Lowicryl HM20 100% |
| -25°C | +20°C | 5°C/h | 9 h (+UV) | Lowicryl HM20 100% |

After embedding in Lowicryl HM20, samples were released from the HPF carriers by dipping the embedded sample in liquid nitrogen. The irregular surface was smoothened by sectioning with a microtome, until the first layer of each sample was on the surface. Sequentially, the resin blocks were immersed in MilliQ (30 min, RT) for rehydration and imaged using a LSM900 upright with Airyscan® 2 detector in SR-4Y mode, to make overviews (10x, C Epiplan-Apochromat 10x, NA 0.4, voxel size: 0.312 × 0.312 × 1.42 μm) and high-resolution z-stacks (40x water-dipping, W Plan-Apochromat 40x, NA 1.0, voxel size: 0.049 × 0.049 × 0.260 μm). In parallel to fluorescence microscopy, reflection microscopy (confocal) was used to visualize the surface of the sample for correlation with the SEM top-view. Reflection images were recorded using the 640 nm excitation laser of the Zeiss LSM900 upright. Images were aligned using the Zen connect module in ZEN 3.7 software (Carl Zeiss Microscopy) to allow for sequential FIB/SEM imaging at the region of interest.

### 3D FIB/SEM

After fluorescence microscopy of the HM20 embedded sample, the sample was coated with a gold sputter coater (Edwards) and transferred to a Zeiss Crossbeam 550 FIB/SEM for volume imaging. Using a top-view, the fluorescence and electron microscopy images were aligned, by overlaying these images in Zen Connect (ZEN 3.7). At the regions of interest, Pt and C were sequentially deposited before a coarse trench was milled using a 30 kV/30 nA FIB probe and smoothened with a 30 kV/3 nA FIB probe. Parameters for serial sectioning imaging were set in Atlas 3D software (Atlas Engine v5.3.3) to allow imaging at a 5×5×5 nm voxel size (FOV 15×20 μm), using a 30 kV/700 pA probe for serial FIB milling. InLens secondary and backscattered (grid -876V) electron microscopy images were simultaneously recorded (line average n=1, dwell time 0.7×4 μs) using an acceleration voltage of 2.0 kV and a probe current of 30 pA. After imaging, the ESB and InLens data were exported from Atlas as a combined file, using a 0.3 ratio of Inlens/ESB.

### Image analysis

Fluorescent images were analyzed using FIJI 1.54f^67^ and ZEN 3.7 (Carl Zeiss Microscopy). Z-stacks were processed by preparing Maximum Intensity Projections (MIP) for quick visualization. Additionally, 3D representations and orthogonal views were used to visualize 3D information. OrientationJ Analysis^68^ and Distribution^69^ were performed on MIPs of 10x z-stacks to visualize cellular and collagen alignment. Overlays of fluorescence and electron microscopy top views were prepared in ZEN 3.7, using Zen Connect, to allow for the alignment of the imaging modalities. After export, FIB/SEM data were denoised in ZEN 3.7 (Total variation, strength 1). The denoised images were then enhanced by local contrast enhancement (CLAHE) in FIJI (block size 49, slope 2.0). Horizontal stripe artifacts were removed after these processing steps, using the FFT power spectrum of the image and invert FFT function in FIJI.

### Chip compression & modelling

Before compression, bonded chips with a PDMS top layer of approximately 200 μm or 800 μm were filled with UV-curable glue (Byllux6308, Byla GmbH) mixed with Oil red O (O9755) for visualization. The filled chips were placed on the bottom part of a custom-made, 3D printed compression device (Supplementary Figure S1). This plastic holder contained one 14.7 N NdFeB magnet (Q-20-05-02-HN, Supermagnete) above which the chip rested with its glass coverslip. Next, the top frame was positioned on top of the PDMS side of the chip so that it could be used to align and support the movable actuator with 5 pins that aligned with the centers of the 5 culture chambers of the chip. The actuator in the top frame contained a second magnet. The distance between the top and bottom part of the compression device was tuned with two screws that extended from the top frame to the bottom frame. After fixation of the chip in the compression device, the stained glue was cured for 30 min using a Honle Bluepoint 2 easy cure (365 nm, 2× 900 s) and left to harden further under compression for 2 days. After disassembly of the compression device, the hardened glue inside the cell culture chambers retained their shapes adopted under compression and the hardened glue in the chips was imaged with confocal fluorescence microscopy from the coverslip side using a LSM900 upright (C Epiplan-Apochromat 10x, NA 0.4, z-step 1.42 μm, averaging 2×) or a LSM900 (10x objective, NA 0.3). The height of the compressed chambers (200 μm PDMS top layer) was measured using an orthogonal view from the 3D microscopy images, which was taken between the pillars. In these images, the height of the fluorescent signal was measured at the thinnest point. The distortion of the pillars under compression (800 μm PDMS top layer) was determined using an orthogonal view from the 3D microscopy images of the pillars, and is expressed as maximum distance of displacement from the uncompressed (straight) situation of the outer pillar surface, facing away from the centre of the chamber.

The magnetic force dependence on the distance between the magnets of the actuation device was measured using a dynamometer. The magnets were separated by a stack of glass slides with different thicknesses and the dynamometer, attached to one of the magnets, was gradually pulled away from the second magnet, until the pulled magnet detached from the stack. The force at the time of detachment was recorded in 5 independent experiments per stack height. The thickness of the stacks, which equals the distances between the magnets, was measured using a caliper to determine the distance between the magnets. The force-distance curve was plotted using the distance data and the force at detachment.

The computational modeling of the compression was performed using Comsol Multiphysics (6.0.0.312). The geometry of one quadrant of the chip and the compressive tip were built (Supplementary Figure S1), because the whole system had 2 symmetrical planes: parallel to XZ-plane and YZ-plane with both planes going through the centered symmetry line. For simulations of the pillar bending, the height of the channel was set to 370 μm (determined from the average heights of all cut open chambers imaged from the side) and ceiling thickness to 800 μm (aimed at ceiling height, controlled by the volume of applied PDMS to the mold). As the PDMS and the column was bonded together, in Comsol simulations, the function “Union” should be used for the geometries on these parts. When selecting the materials for the components, the tip was assumed to be non-deformable (built-in material from Comsol library: steel AISI 4340) and the properties of the PDMS were set according to previously published characteristics (density: 1065 kg/m^3,70^ Young’s modulus: 1.5 MPa,^71,72^ Poisson’s ratio: 0.495^73^). To set up the solid mechanics, the *hyperelastic node* was added, and the Mooney-Rivlin-5-parameter model was selected with the following parameter values in MPa: C10=-0.46701; C01=0.88163; C11=-3.91278; C20=1.54142; C02=3.11667 with a bulk modulus of 3 GPa.^74^ *The contact node* was used with these contact pairs: tip’s surface and the top of the PDMS layer, as well as the top and bottom surfaces of the chamber. The augmented Lagrangian method was chosen over the penalty method. *Symmetry nodes* were defined at these 2 planes: parallel to XZ-plane and YZ-plane. The *fixed constraint node* was assigned to the side-faces of the glass slide. The last node to be added was *boundary displacement* with the function z=-x assigned to the surface of the tip. The meshes were created with the focus on the contact surfaces between the PDMS block, the tip, and the pillar whereas the other components could be treated with reduced coarser meshes. The solver chosen was auxiliary sweep from 0 to 1 with a step size of 0.1 for the parameter x. When x=0.2, the tip was in the first contact the PDMS surface; when x=1, the tip had travelled 0.26 (mm). Point evaluation with “solid.disp (mm)” was used to evaluate the displacement in mm of the following points: the tip (point 47), the top of the ceiling (point 42), the bottom of the ceiling (point 41), six points on the bottom half of the pillar (points 16 to 11, shown as points 1-6 in Supplementary Figure S1). Surface evaluation with “solid.tnz (N)” was used to evaluate the contact force, and the top surface of the ceiling was chosen.

## Author contributions

JS: Conceptualization, Data Curation, Investigation, Methodology, Visualization, Writing – original draft, Writing – review & editing. DW: Conceptualization, Formal analysis, Investigation, Methodology, Writing – review & editing. MEC: Investigation, Methodology, Writing – review & editing. MKL: Investigation, – review & editing. RvdM: Investigation, Methodology, Writing – review & editing. RR: Investigation, Writing – review & editing. ND: Investigation, Writing – review & editing. MPL: Formal analysis, Investigation, Methodology. JR: Formal analysis, Methodology. WHN: Resources. AvdM: Conceptualization, Funding acquisition, Methodology, Writing – review & editing. PJ: Conceptualization, Funding acquisition, Methodology, Supervision, Writing – review & editing. NS: Conceptualization, Funding acquisition, Methodology, Supervision, Writing – original draft, Writing – review & editing. AA: Conceptualization, Funding acquisition, Methodology, Supervision, Visualization, Writing – original draft, Writing – review & editing.

## Data availability

Data presented in this article, including imaging files with metadata, image analysis (orientation and compression) and spectroscopy data are available at Radboud Data Repository at https://doi.org/10.34973/zsjf-926”.

## Acknowledgements

The authors thank Harrie Weinans and Ralph Sakkers from University Medical Center Utrecht for providing the hMSCs. The authors also would like to thank Tom Boonen and Carla Cofiño-Fabres from the University of Twente for their help during the design phase of our device and for sharing their knowledge on OoC cell cultures in these types of devices, José Manuel Rivera Arbeleaz from the University of Twente for help with the modeling of the mechanical actuation, and Marcel de Bruine from University of Twente for 3D printing of the actuator parts.

