## Supplementary Figures & Supplementary Table for "A workflow integrating organ-on-chip culture and correlative 3D light and electron microscopy for microtissue analysis"

**Supplementary Material**

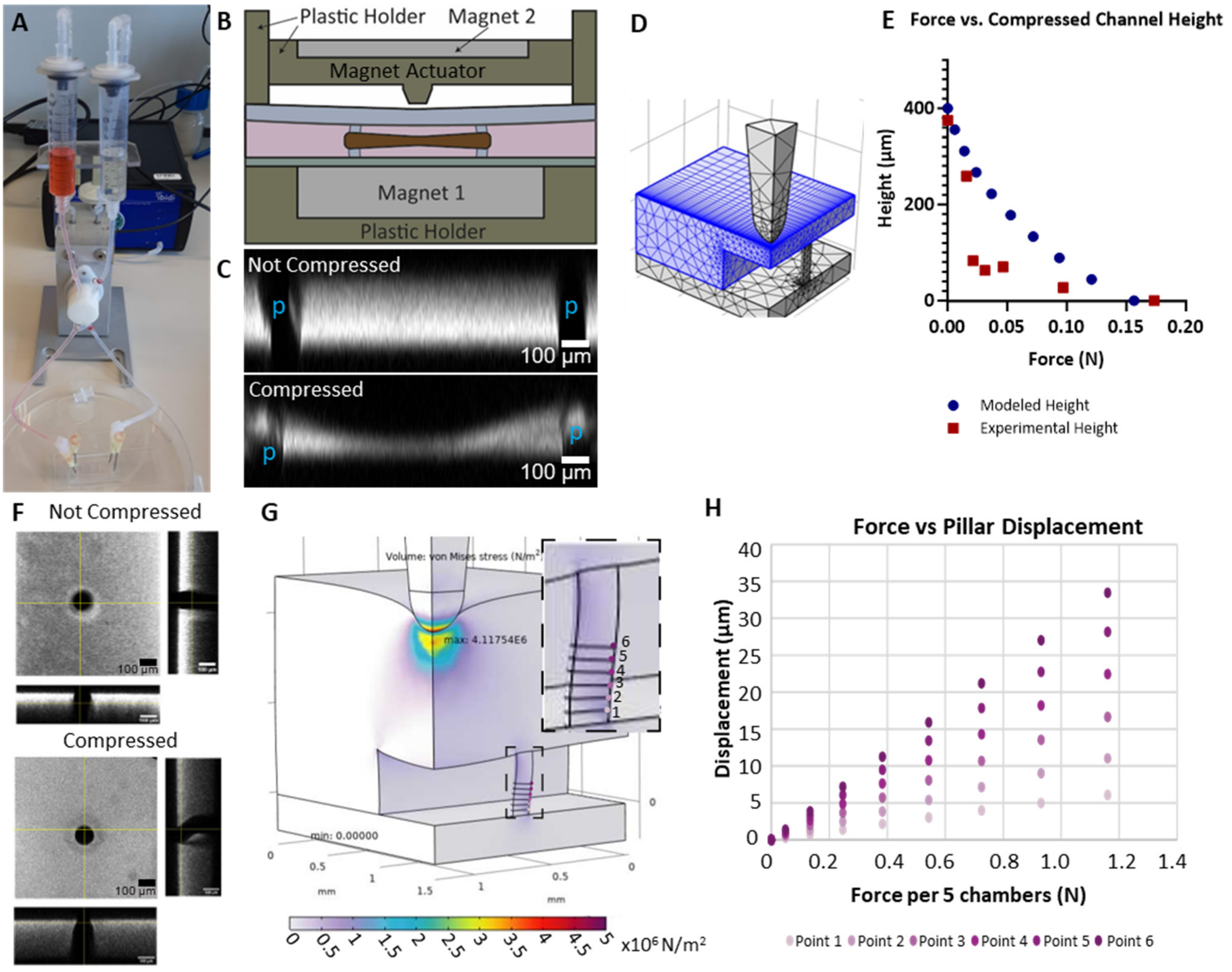


**Supplementary Figure S1**

*A) Representation of our organ-on-a-chip platform connected to the Ibidi® flow system including 2 syringes as medium reservoir. The chip rests with the coverglass on the 3D printed bottom holder containing the first of the two 14.7 N NdFeB magnets. A 3D printed frame is used to align and stabilize the second magnet, which rests inside the 3D printed actuator with one pin compressing the PDMS-ceiling of each culture chamber. The distance between the magnets is variable to enable the use of different forces of compression. B) Schematic representation of compression device based on magnetic forces and an actuator tip. C) Side-view of fluorescent images from Oil-red-O stained UV-curable glue, loaded on chip and polymerized during no compression (top) or compression (bottom). Black regions indicated with a blue "p" indicate the pillar locations. D) Representation of one quadrant of the chip, as reconstructed in Comsol Multiphysics. The PDMS (in blue, 200 µm top layer), compressive tip (grey top) and bottom glass slide (grey bottom) were included in the model. E) Modeled (blue) and experimentally measured (red) height of the chambers in our chip upon magnetic compression at various magnetic forces in chips with 200 µm top layers of PDMS. F) Fluorescence microscopy images and corresponding orthogonal views (at crosshair) of one pillar of chips filled with UV-curable glue mixed with a Oil-red-O staining (top layer PDMS: 800 µm). Chips were imaged after curing the glue at 365 nm without (top) and during (bottom) magnetic compression (0.49 N on all 5 chambers). Scalebars: 100 µm. G) Simulations of magnetic compression on culture chambers with a top PDMS layer of 800 µm. Model of one quarter of a culture chamber under compression by the pin of the magnetic actuator. The outwards displacement of points 1-6 at the outer surface of the pillars (as indicated in E) was modeled. The color scale indicates the magnitude of von Mises stresses under compression of 0.098 N (one fifth of 0.49 N for only 1 chamber). H) Computationally modeled pillar displacement under magnetic compression, as described in F. Pillar displacement of points 1-6, as indicated in C, was plotted against the applied force. The applied force is given as total force applied to the complete chip with 5 parallel chambers.*

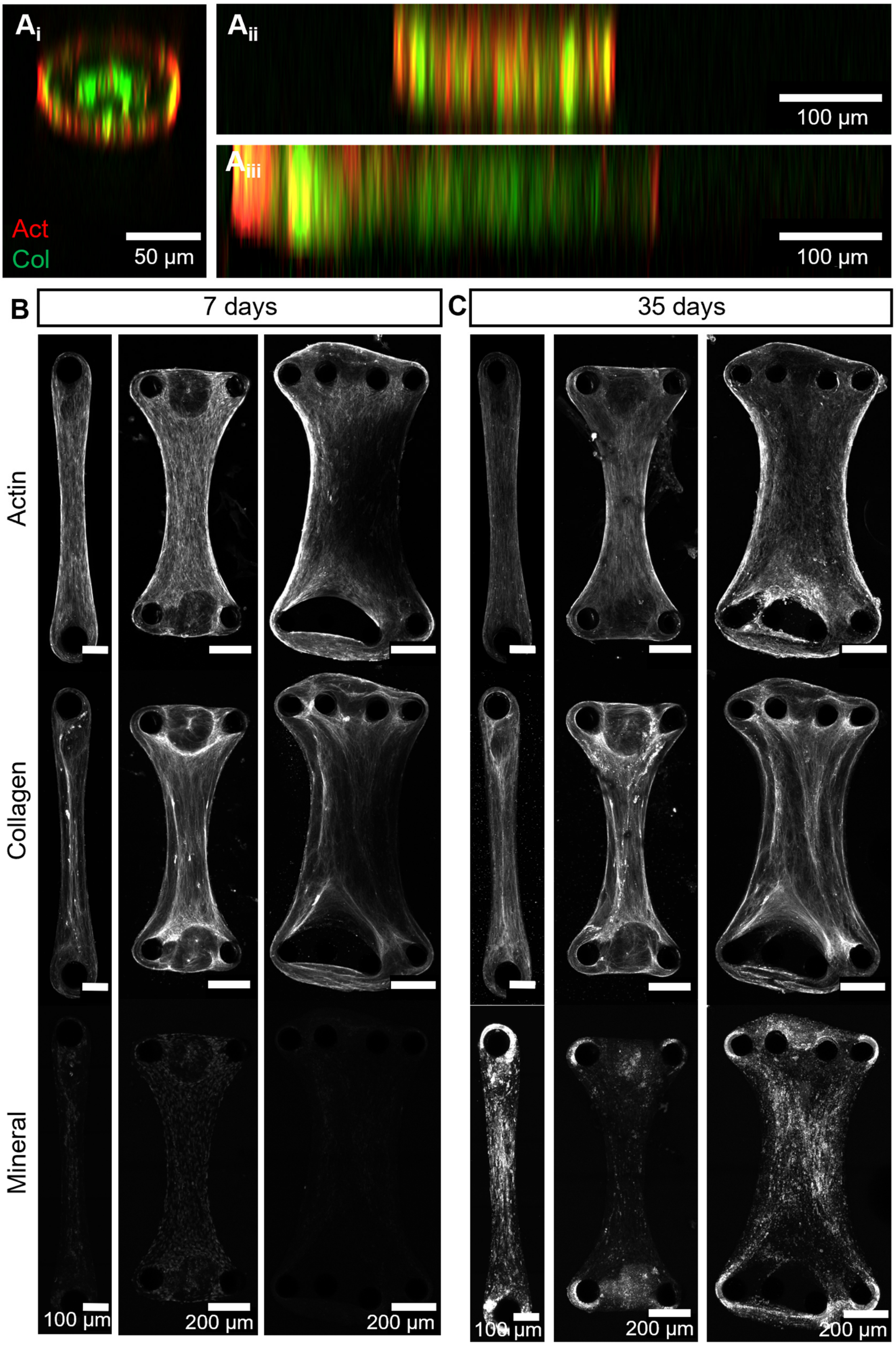


**Supplementary Figure S2**

*A) Orthogonal views of microtissues after 7 days of differentiation in chips with varying designs showed the dependence of the culture structure on the orientation of the pillars. In chips with two single pillars ($A_i$) the cultures showed a symmetrical, round to oval shape, while in chips with pillar arrays of two ($A_{ii}$) or four ($A_{iii}$) pillars, the culture showed a flatter and stretched morphology. (Actin: SiR-actin, red; Collagen: CNA35-OG®488, green). B) Maximum intensity projections of single channels from the multicolor images shown in Figure 2A, after 7 days of differentiation. C) Maximum intensity projections of single channels from the multicolor images shown in Figure 2B, after 35 days of differentiation.*

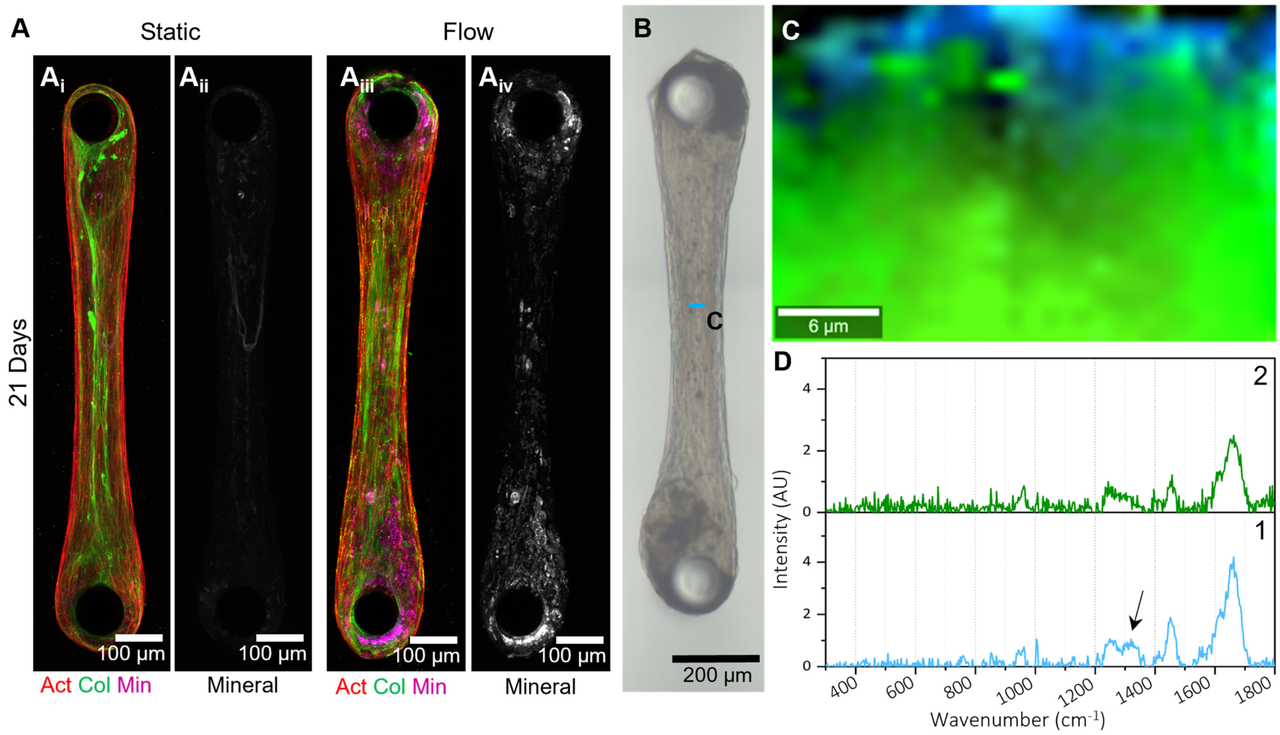


**Supplementary Figure S3**

*A) Fluorescently stained (Actin: SiR-actin, red, Collagen: CNA35-OG®488, green; Mineral: Calcein blue, magenta) static ($A_i$) bone-on-a-chip culture after 21 days of differentiation showed no visible mineral signal yet. Isolated mineral signal in $A_{ii}$. Flow exposed (0.8 ml/min, $A_{iii}$) bone-on-a-chip culture after 21 days of differentiation showed the presence of mineral. Isolated mineral signal in $A_{iv}$. B-D) Live Raman mapping on-a-chip. B) Optical image of a living bone-on-a-chip culture exposed to fluid flow (0.8 ml/min) after 21 days of differentiation with an indicated region of interest in which Raman depth mapping was performed. C) Spatial distribution of TCA components identified in Raman depth map localized in the center of the bone-on-a-chip culture. D) Spectra matching with the components in C identified 1) cellular components (blue) and 2) matrix (green) with a low presence of mineral.*

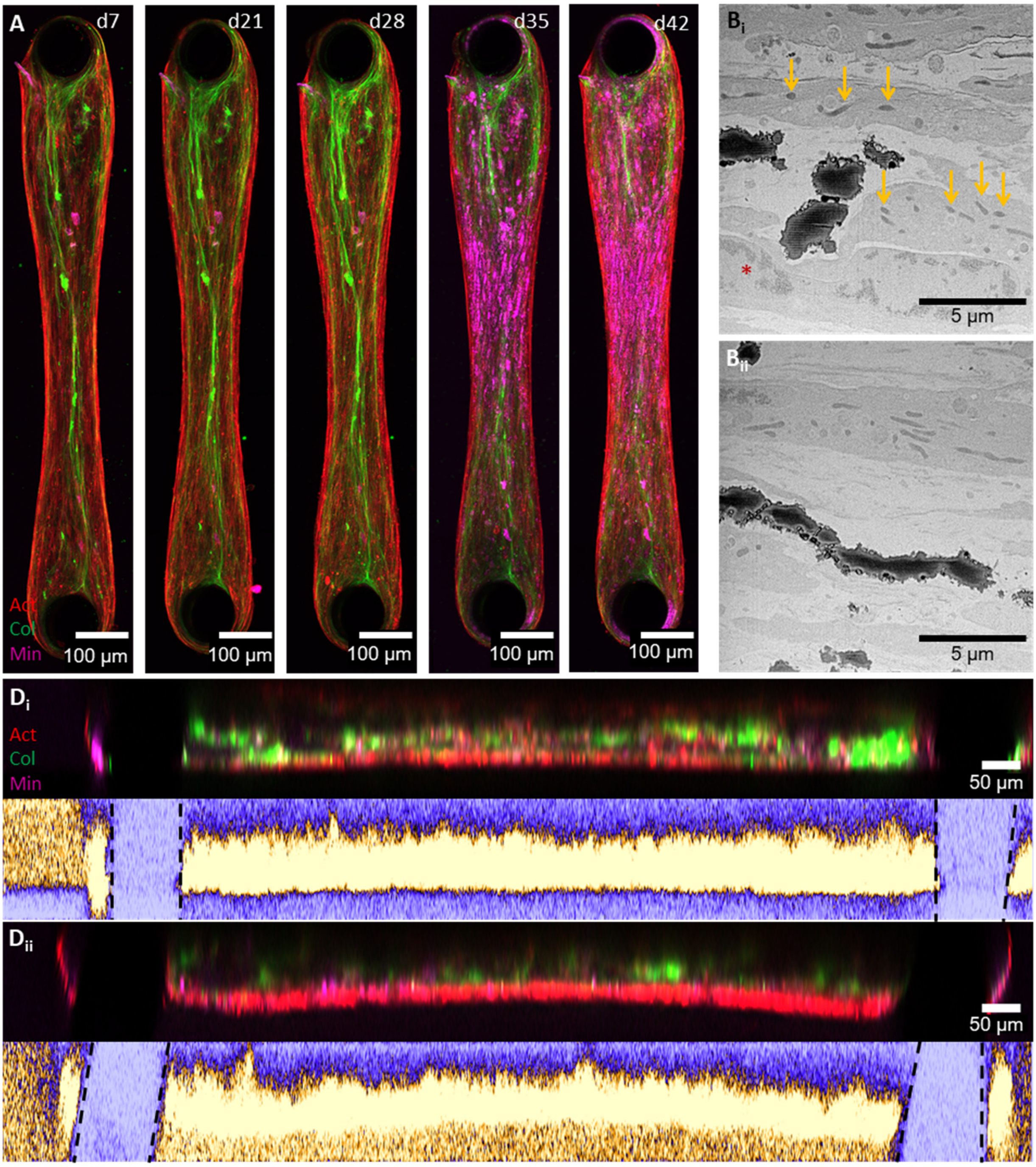


**Supplementary Figure S4**

*A) Live fluorescence imaging over time to monitor cellular organization (Actin: SiR-actin, red), collagen deposition (CNA35-OG®488, green) and mineralization (Calcein Blue, magenta) showed increasing mineralization over time, starting at day 35. (Images at different timepoints from the same culture as shown in Figure 4A and Figure 3A-C). B) Single images from 3D FIB/SEM stack recorded after 42 days of differentiation, showed the presence of cells with organelles ($B_i$: mitochondria →, nucleus *), with collagen and mineral between the cells. The amount of collagen increases while going deeper into the culture ($B_i$ = 5 μm, $B_{ii}$ = 11.57 μm). D) Comparison of pillar alignment in the resliced image (XZ) of a culture showing no twist ($D_i$) or a twist ($D_{ii}$) at day 35. The top image shows the resliced image with all three channels (Actin: SiR-actin, red; collagen CNA35-OG®488, green; mineral, Calcein Blue, magenta). The bottom image shows the mineral channel with ICA LUT to indicate the position of the culture (yellow) and pillars (purple).*

**Supplementary Table S1 - Sample Number per Chip Design (Fluorescence Microscopy)**

| Design | # Microtissues (Total) | # Chips | # Experiments |
|---|---|---|---|
| 4 pillar design | 4 | 1 | 1 |
| 2 pillar design | 3 | 1 | 1 |
| 1 pillar design - Mineralization* | 7 | 2 | 2 |
| 1 pillar design - Actin Orientation* | 8 | 4 | 3 |

* For mineralization experiments, only samples stained with SiR-Actin, CNA-OG®488 and calcein blue were included. For actin orientation experiments all samples that were stained with SiR-Actin were included. 5 samples were used for both analyses. Samples in which the pattern development could not be clearly attributed to one group (e.g. very minor change or increase only observed at one timepoint, but not at a later timepoint) were excluded from the actin orientation analysis.
